# Corticospinal excitability at rest outside of a task does not differ from task intertrial intervals in healthy adults

**DOI:** 10.1101/2024.02.05.578973

**Authors:** Kate Bakken, Chris Horton, Mitchell Fisher, Corey G. Wadsley, Ian Greenhouse

## Abstract

Human corticospinal excitability modulates during movement, when muscles are active, but also at rest, when muscles are not active. These changes in resting motor system excitability can be transient or longer lasting. Evidence from transcranial magnetic stimulation (TMS) studies suggests even relatively short periods of motor learning on the order of minutes can have lasting effects on resting corticospinal excitability. Whether individuals are able to return corticospinal excitability to out-of-task resting levels during the intertrial intervals of behavioral tasks that do not include an intended motor learning component is an important question. Here, in twenty-six healthy young adults, we used single-pulse TMS and electromyography (EMG) to measure motor evoked potentials (MEPs) during two different resting contexts: 1) intertrial intervals of a choice-reaction time task, and 2) outside the task. In both contexts, five TMS intensities were used to evaluate possible differences in recruitment of corticospinal output. We hypothesized resting state excitability would be greater during intertrial intervals than out-of-task rest, reflected in larger MEP amplitudes. Contrary to our hypothesis, we observed no significant difference in MEP amplitudes between out-of-task rest and in-task intertrial intervals, and instead found evidence of equivalence, indicating that humans are able to return to a stable motor resting state within seconds after a response. These data support the interpretation that rest is a uniform motor state in the healthy nervous system. In the future, our data may be a useful reference for motor disorder populations with an impaired ability to return to rest.

## Introduction

Motor system excitability modulates dramatically during behavior. A considerable amount of research has characterized such modulation before, during, and after motor responses. However, few studies have directly compared motor excitability during in-task intertrial intervals with out-of-task rest contexts. Both in-task intertrial intervals (structured) and out-of-task (unstructured) rest are contexts in which volitional motor activity is absent, but whether corticospinal excitability is consistent between these contexts is not known. Comparisons of ‘rest’ are important because numerous behavioral studies use intertrial intervals as a baseline reference for determining the effects of in-task behavioral manipulations. Moreover, whether individuals are capable of returning to a stable state, equivalent to out-of-task rest, between epochs of movement may be important for characterizing healthy motor system function.

Corticospinal excitability (CSE) can be measured with transcranial magnetic stimulation (TMS) over primary motor cortex (M1), and TMS studies of CSE in the context of motor tasks have examined plasticity induced through motor learning over the span of hours and days (for reviews see Bestmann & Krakauer 2015; Carson et al. 2016; Spampinato et al. 2013). Effects of motor learning on corticospinal activity have also been observed across shorter timescales. For example, repeating thumb movements along a fixed trajectory for a period of as little as five minutes results in TMS-elicited thumb movements along the trained trajectory, providing evidence for short-term motor learning and the potential for plasticity to occur rapidly in the brain (Classen et al. 1998; Bütefisch et al. 2000; Suleiman et al. 2023). Modulation of CSE can also occur without any explicit motor learning, such as immediately before voluntary movement or immediately before a state of voluntary relaxation (Bestmann & Duque 2016; Duque et al. 2017; Chen & Hallett 1999; Suzuki et. al. 2015). Evidence of transient motor system modulation calls into question the capacity of the motor system to return to a stable resting state between trials of a behavioral task, and thus, whether intertrial measures can serve as a valid baseline reference.

Changes in CSE during movement preparation extend beyond muscles involved in responding. For example, TMS-elicited motor evoked potential (MEP) amplitudes decrease during the preparatory period of a delayed response task in task-relevant and task-irrelevant muscles (Duque et al. 2009, 2010, 2012, 2017; Greenhouse et al. 2015; Lebon et al. 2019; Labruna et al. 2021; Hannah et al. 2020). The potential influence on intertrial rest of these presumed transient changes to CSE is under-researched. Intertrial dynamics may hold valuable information regarding mechanisms responsible for modulating motor system excitability. Establishing whether healthy individuals return to a stable resting baseline level of CSE could aid future investigations in clinical populations. Moreover, examining a range of TMS intensities may provide more complete information about the input-output state of the corticospinal pathway since differences can manifest within separate segments of an MEP recruitment curve. For example, sensitivity to weaker stimulation intensities may increase shortly after the execution of a motor response while the maximum attainable MEP amplitude may remain stable. Such non-linear adjustments in input-output relationships may point to the lingering influence of specific mechanisms within the corticospinal pathway after a response is executed.

Here, we used TMS to compare resting CSE between out-of-task and in-task contexts. Input-output MEP recruitment curves were measured using five TMS intensities during out-of-task rest and within intertrial intervals (ITIs) of two separate motor response tasks. All measurements were taken from the left first dorsal interosseous (FDI) muscle, with in-task measurements taken when the FDI was both task-relevant and task-irrelevant. We hypothesized that the recruitment curves of individuals at rest between trials of a task would exhibit enhanced CSE compared with those produced for the same individual outside of a task. Specifically, we predicted an overall leftward shift in the recruitment curve and a steeper slope, arising from lingering activity associated with task responses. Such a result would be consistent with the interpretation that rest is not a uniform state inside and outside a task context. Alternatively, no differences in the recruitment curves across conditions would indicate a stable, uniform resting state of motor excitability. This result would support the use of intertrial rest as a baseline reference for comparisons with non-resting behavioral states.

## Methods

### Participants

Twenty-six young, neurologically healthy, right-handed adults volunteered to participate in this study (16 female, 10 male; mean age of 20.1 ± 3.1 years old). Data from one participant were excluded from the analysis due to high levels of background EMG activity during the task. Written informed consent was obtained from each participant before data collection and all participants completed the entire study. All participants were screened for contraindications prior to TMS, including family or personal history of seizure and personal history of head trauma or fainting. The institutional review board of the University of Oregon approved this study.

### Experimental Design

Data were collected during three experimental conditions: out-of-task rest, a task involving a choice between the two responding hands (hand-choice), and a task involving a choice between two responding fingers of the right hand (finger-choice). These three conditions were completed in one data-collection session for 22 of the 26 participants, and two separate sessions for the other four participants due to time constraints. For all participants, the out-of-task rest condition was completed first, and the order of the hand-choice and finger-choice conditions was counterbalanced across participants. For all conditions, the participants were asked to sit in a relaxed position approximately two feet in front of a computer monitor, with their forearms and hands resting flat on a table in front of them. Participation lasted approximately 2.5 hours in total.

### Out-of-task EMG and TMS

Electromyography (EMG) was collected over the left FDI to record MEPs and the C7 vertebra to detect TMS artifacts, with a ground electrode positioned over the left head of the ulna (Figure 1). Additional electrodes were placed over the FDI and abductor digiti minimi of the right hand for measuring response-related EMG bursts, but these data are not discussed here. The skin under each electrode was lightly exfoliated and cleaned with alcohol prior to affixing bipolar Ag/Cl electrodes to the surface of the skin. All electrode channels were connected to a Bagnoli (Delsys) EMG amplifier (1000x) and sampler (5000 Hz; bandpass filter 50-450 Hz). EMG data were recorded onto a computer using the VETA toolbox (Jackson & Greenhouse, 2020) in MATLAB (Mathworks, Natick, MA) throughout each task. Participants were shown their online EMG recordings and were instructed to minimize activity in all channels when not responding.

**Figure 1.**
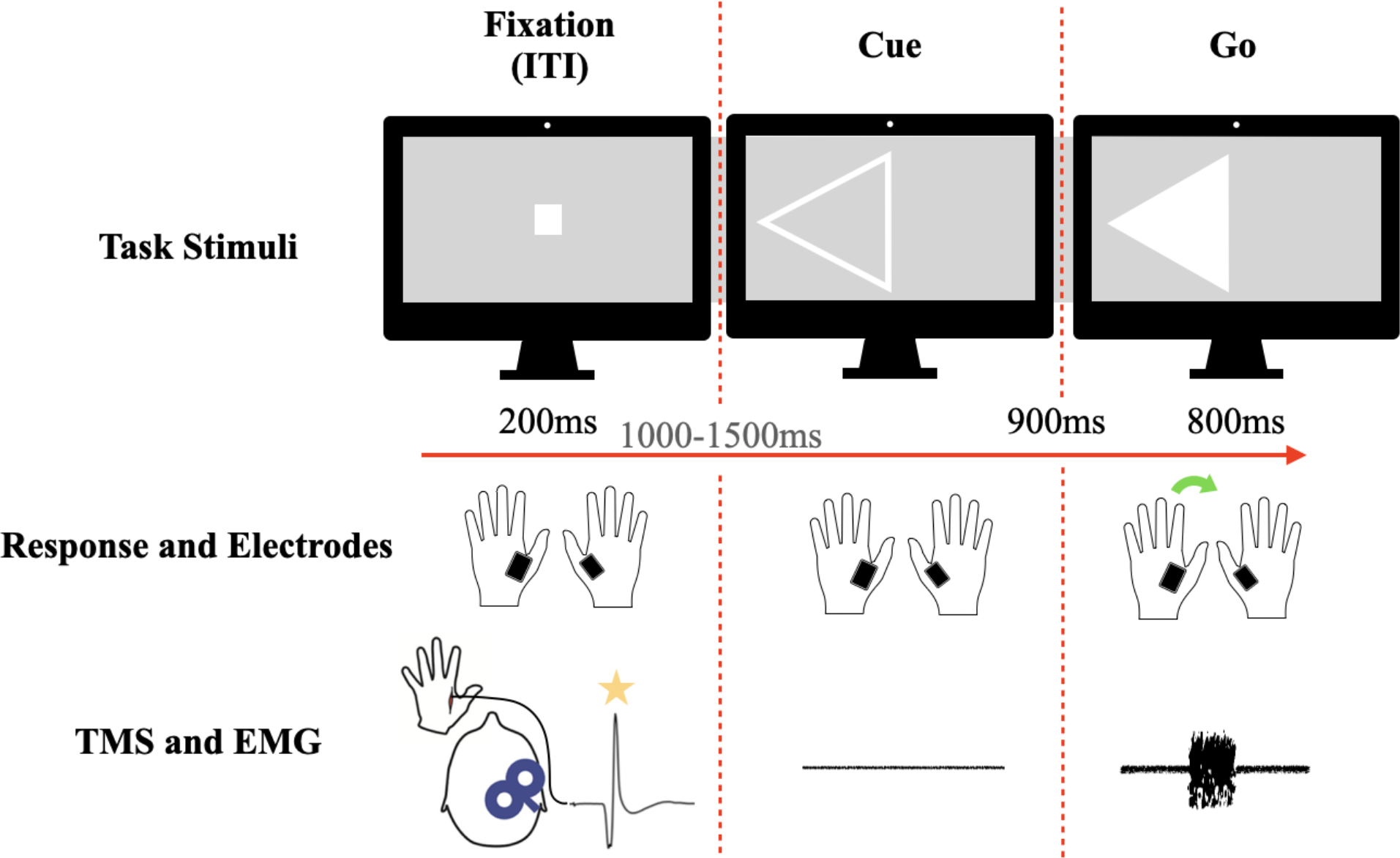
Task stimuli consisted of left and right-pointing arrows. One arrow became bold to cue the participant to prepare a corresponding response. The arrow then filled in to signal the execution of the prepared response. Participants responded to left and right arrows using the left and right index fingers, respectively, during the hand-choice task (depicted) with surface EMG recorded from the responding first dorsal interosseous (FDI) muscles. The right FDI and ADM served as the responding muscles in the finger choice task (not shown). TMS administered over the right M1 elicited MEPs (star) in the left FDI at fixation onset to provide measurements of resting CSE during the task. An example EMG trace is shown from a single trial.

TMS hot-spotting and thresholding were performed to reliably elicit MEPs from the left FDI. The TMS coil was first positioned 2 cm anterior and 5 cm to the right of the vertex of the head and angled approximately 45º off the midline to produce a posterior-anterior flowing current over M1. The coil was repositioned in increments of approximately 1 cm until reliable MEPs were observed in the left FDI EMG recording. To determine resting motor threshold (RMT), TMS intensity was initially set at 30% maximum stimulator output (MSO) and was adjusted by 2% MSO until MEPs with amplitudes above 0.5mV were visualized on 5 out of 10 consecutive pulses over the motor hotspot.

Out-of-task rest measurements consisted of 70 TMS pulses over the left FDI hotspot in right M1 at 90, 110, 130, 150, and 170% of the participant’s RMT. The order of TMS intensities was randomized across measurements with 14 pulses at each intensity. The participant was instructed to sit with their body as relaxed as possible, with reminders provided as needed by the experimenter. EMG activity was visualized online by the experimenter on a screen adjacent to the participant and recorded with the VETA toolbox.

### Behavioral Tasks

The two behavioral tasks used the same behavioral stimuli and event timing and only differed in the response configuration. Both tasks consisted of three epochs: Fixation, Cue, and Go (Figure 1). In the fixation epoch, a centered white rectangle (20 x 20 pixels) appeared 300 ms after trial onset for 200 ms. The cue epoch started 1000 to 1500 ms after trial onset (randomly drawn from a uniform distribution) and cued the participant to prepare either the left or right response (hollow leftward or rightward pointed triangle, 140 x 150 pixels). The go epoch (triangle filling white) began 900 ms after cue onset and stayed on the screen for 800 ms or until a button press was detected. Each trial was followed by an intertrial interval (blank gray background) which lasted until 7.6 s had elapsed from trial onset. The participant was instructed to relax during the fixation period, to prepare but not execute the forthcoming response during the cue epoch, and to execute the cued response as fast as possible during the go epoch. Both tasks consisted of four blocks, which alternated between 59 and 60 trials each, for a total of 238 trials. Of these 238 trials, 14 were ‘catch trials,’ in which the go stimulus never appeared to discourage participants from responding prematurely.

Behavioral data were collected via button presses from custom-built response boards (Makey Makey v.1.2; Joylabs). During the finger-choice task, participants responded to the left go stimulus by pressing a button using a lateral movement of the right index finger and responded to the right go stimulus by pressing a button using a downward movement of the right pinky finger. During the hand-choice task participants responded to the right go stimulus with the right index finger and the left go stimulus with the left index finger. In this case, lateral movements of the index fingers were used to press buttons on the sides of a box positioned beneath the stimulus presentation display. Otherwise, all aspects of the tasks were identical. After the researchers explained each task, participants completed a short practice block until they expressed readiness to begin testing, approximately 10 trials.

### In-task TMS

In-task TMS was administered at the same intensities as out-of-task TMS (90, 110, 130, 150, or 170%) and equally distributed across the cued left and right response directions. TMS was administered on 210 trials for both the finger-choice and hand-choice tasks (14 pulses per TMS intensity level at the fixation onset (ITI) and 14 pulses per intensity level, per hand 800ms into the cue epoch).

Here, we analyze in-task TMS pulses administered at fixation onset as this pulse time was at the end of the ITI and serves as a measure of resting CSE in the context of a motor task. Similar to the out-of-task context, online EMG was monitored during the task to ensure participants remained at rest when not responding, where the experimenter instructed the participant to relax if increased EMG activity was noted outside of go epochs.

### Data Analysis

All surface EMG data were pre-processed and visualized using the VETA toolbox in MATLAB using a two-step procedure. First, MEP and EMG events were automatically detected using the ‘findEMG.m’ function, and then all the data were visualized using the ‘visualizeEMG.m’ function and associated graphical interface. The three conditions (rest, finger-choice, hand-choice) were visualized separately. Trials that included excessive EMG activity and/or EMG activity overlapping with MEPs or MEPs detected at the wrong intervals were excluded from the analysis.

Mean MEP amplitudes and root mean square (RMS) of the EMG activity during the 100 ms preceding TMS pulses were calculated for each TMS intensity for the out-of-task, hand-choice ITI, and finger-choice ITI conditions. A Boltzmann function was fitted to the mean MEP amplitudes across the five TMS intensities for each of the conditions (Kukke et. al. 2014) using the ‘sigm_fit.m’ function in Matlab:

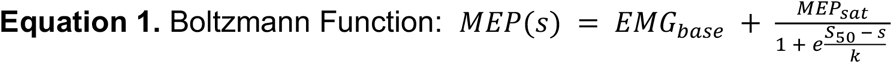

where *MEP*_*sat*_ is the plateau value at high stimulation intensities, *EMG*_*base*_ refers to the participant’s EMG baseline at rest, ^*S*^50 is the stimulation intensity that produces a MEP halfway between *EMG*_*base*_ and *MEP*_*sat*_ and *k* is the change in stimulus intensity from ^*S*^50 that relates to a 73% change in a participant’s *MEP(s)* (Kukke et. al., 2014).

The slope of the line of best fit for each curve was calculated using the ‘sigm_fit_val.m’ function in Matlab, and maximum slope was determined.

### Statistics

Statistical tests were analyzed with Bayesian repeated-measures analysis of variance (ANOVA) using the program JASP (JASP team 2023). All models included random slopes for all repeated measures factors and were fitted across x100,000 iterations with participant modeled as a random intercept (van den Bergh et al. 2022). Normality of data and model-averaged residual plots were checked in JASP using a Q-Q plot of residuals. No transformations were required, as there was no non-normal data. Evidence for main effects and interactions were determined using Bayes factor in favor of the alternative hypothesis (BF_10_ ± percent error), where values greater than 1 indicate support for the alternative hypothesis and values less than 1 support the null hypothesis. The strength of evidence was determined using a standard BF_10_ classification table (BF_10_ < 0.3: moderate evidence for the null hypothesis; 0.3 ≤ BF_10_ ≤ 3: weak evidence for the null hypothesis; BF_10_ > 3: moderate evidence for the alternative hypothesis; van Doorn et al. 2021). Main effects and interactions were further evaluated using post-hoc pairwise comparisons that were performed using Bayesian paired t-tests in JASP. The null hypothesis was zero difference across conditions, with the alternative being a difference not equal to zero. A Cauchy prior distribution was assumed for the null. All data are presented as non-transformed mean ± standard deviation.

Mean RMS of EMG activity from -60 to -10 ms preceding the TMS pulses and mean MEP amplitudes were assessed with 2-way repeated-measures ANOVA using the factors Condition (rest, hand-choice ITI, finger-choice ITI) and TMS Intensity (90, 110, 130, 150, 170% RMT) to test our hypothesis that MEP amplitudes would increase during inter-trial intervals of the finger and hand epochs compared to the out-of-task resting state.

## Results

Participants performed the motor tasks correctly, as indicated by the high levels of accuracy across the two tasks (finger: 98.65 ± 3.88% correct; hand: 96.81 ± 6.15% correct). Three participants were excluded from the hand-choice behavioral data analysis due to a malfunction of the button response device. Overall accuracies indicate that participants were behaving as expected and executing trial-wise responses. We did not evaluate other behavioral metrics as we had no hypotheses about relationships between resting CSE values and task performance.

Participants had an average RMT of 43 ± 7% MSO (all < 65% MSO). For the 2-way repeated-measures ANOVA on RMS of EMG -60 to -10 ms from TMS onset, there was no effect of Condition (p = 0.13) and no effect of TMS intensity (p = 0.07). A Greenhouse-Geisser sphericity correction was made in calculating these effects, as Mauchly’s test of sphericity indicated the assumption of sphericity was violated (p < 0.05) when the uncorrected repeated-measures ANOVA was initially run. This indicates participants did not have differences in background EMG activity immediately preceding the TMS pulse.

For the 2-way repeated-measures ANOVA on MEP amplitudes, there was strong evidence for a null effect of Condition (BF_10_ = 0.259), with no difference between out-of-task and hand-choice (0.146 mV, posterior odds = 0.086), between out-of-task and finger-choice (0.381 mV, posterior odds = 0.224), or between the two task conditions (1.463 mV, posterior odds = 0.224). There was strong evidence for an effect of TMS intensity (BF_10_ > 10), where MEP amplitude increased reliably across intensities (all post-hoc pairwise comparisons BF_10_ >1). There was very low evidence for an interaction between Condition and Intensity (BF_10_ = 0.112).

The recruitment curves for each condition (rest, hand, finger) at each intensity (90, 110, 130, 150, 170% RMT), with mean MEP amplitude (mV) ± sem, are presented in Figure 2. The group mean (std) of the maximum recruitment curve slope values were 0.055 (0.035), 0.060 (0.028), and 0.051 (0.027) for the rest, hand, and finger conditions, respectively. Statistical tests comparing maximum slope across conditions were not appropriate because the model fitting approach depends on within-subject variance that cannot be recycled for group-level comparisons. However, these slope values are highly similar, and visual inspection indicated overlapping distributions.

**Figure 2.**
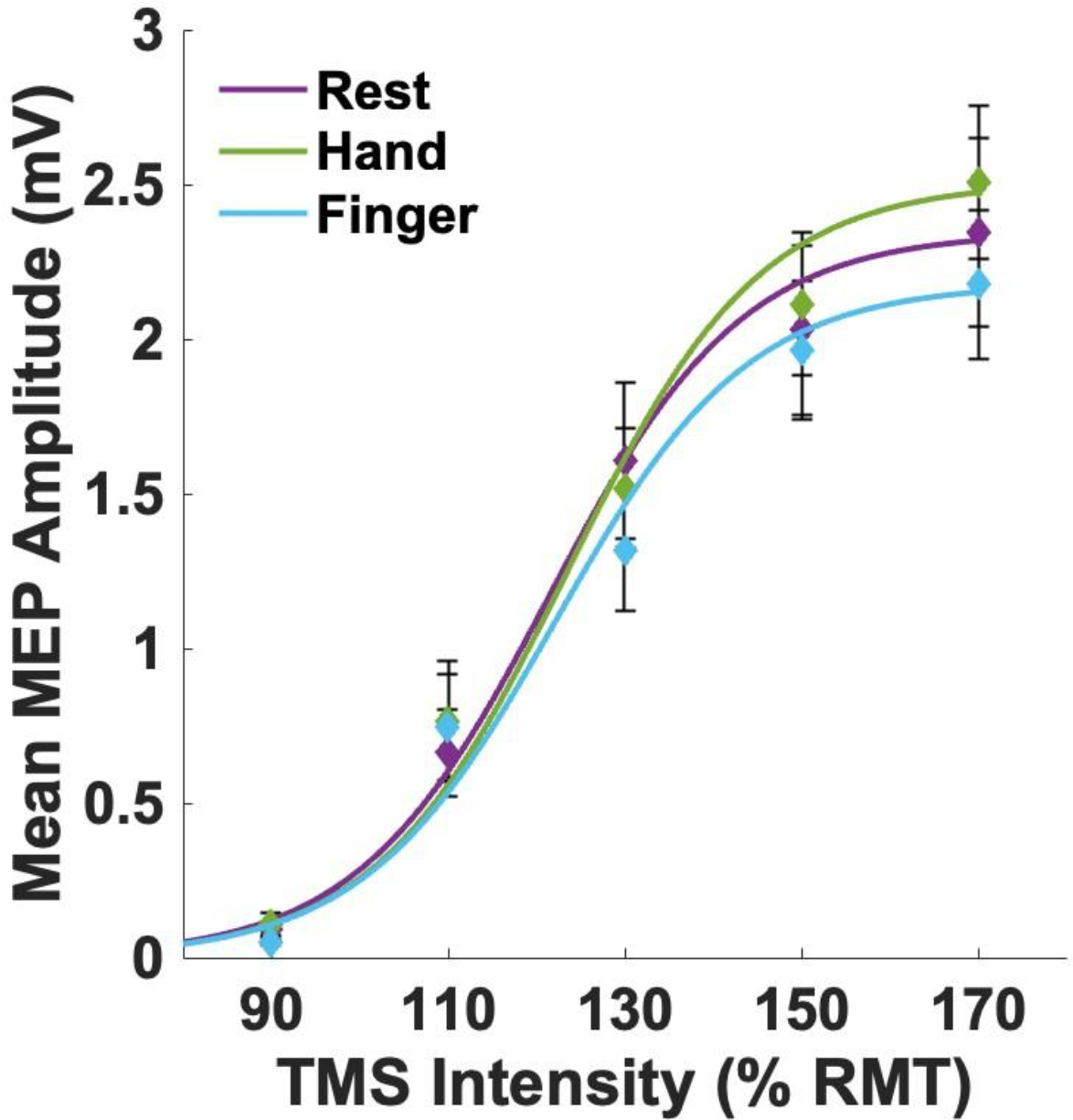
Mean MEP amplitudes (mV) across six TMS intensities (0, 90, 110, 130, 150, 170 % RMT) for each of the 3 conditions (Rest, Hand, Finger) were fit with a Boltzmann function (solid lines) to derive recruitment curves. Mean ± sem MEP amplitude (diamonds) increased with increasing %RMT but did not differ significantly across conditions

Together, these results indicate the distribution of MEP amplitudes across TMS intensities is highly overlapping between out-of-task and ITI rest. Thus, resting CSE did not change in the context of the task.

## Discussion

The present study compared single-pulse TMS measurements of CSE across a range of stimulation intensities between rest outside a task and during the inter-trial intervals of instructed-delay two-choice reaction time tasks. Our hypothesis that MEP amplitudes would increase during inter-trial intervals compared to the out-of-task resting state was not supported. Instead, we observed converging evidence in favor of the null hypothesis that MEP amplitudes from the different resting conditions are equivalent across a range of TMS intensities. Moreover, recruitment curve slopes calculated for each condition were in a similar range for the three conditions. Overall, the results indicate inter-trial CSE returns to an out-of-task rest state during the performance of delayed response tasks.

Many TMS experiments depend on inter-trial measurements as a reference for CSE modulation during phases of a behavioral task. Our current data indicate in healthy participants that the baseline measurements of CSE inside and outside of a behavioral task context are similar regardless of whether MEPs are measured from responding or non-responding hand muscles. This suggests the human corticospinal pathway can return to a consistent resting state within a matter of seconds after the execution of a response, and the inter-trial baseline may serve as a reliable proxy for out-of-task resting CSE measurements. The current data extend previous studies that have compared out-of-task and within-task resting MEP measurements at a single TMS intensity (e.g. Wadsley et al. 2023; Greenhouse et al. 2015) by accounting for the possible influence of variable TMS intensities.

Our exploratory analysis of maximum recruitment curve slopes across participants and conditions adds to the growing research on input-output relationships of the corticospinal pathway. The slope of the line of best fit for the recruitment curve represents the sensitivity of the motor system to increasing TMS intensities, with higher slopes indicating greater sensitivity to increasing cortical input (Kukke et al. 2014). Our results support the application of the Boltzmann function at both the group and individual levels for fitting these data. The similarity in the estimated curve slopes for the different conditions, in the context of the omnibus ANOVA results for mean MEP amplitudes, lends further support to the interpretation that rest is a uniform state inside and outside the context of the delayed response tasks.

The lack of CSE modulation during within-task rest is somewhat unexpected. Previous studies using behavioral tasks similar to ours, i.e. involving repeated finger responses across trials, have established changes in short-term CS plasticity measured as adjustments in TMS-elicited movement trajectories (Classen et al. 1998; Bütefisch et al. 2000; Suleiman et al. 2023). Such plasticity suggests repeating finger movements changes the state of the CS pathway over the span of minutes, however, we did not observe evidence for CSE changes in the responding effector during the ITI. Differences from this previous work may be explained by features of our chosen response tasks. Specifically, in those studies movements were repeated in consecutive trials whereas here, participants chose between two response options on a trial-by-trial basis. Alternatively, it is possible MEP amplitudes are more dynamic than TMS-elicited movement trajectories. MEP amplitudes change dynamically from the ITI during the response preparation period (Duque et al. 2017), but less is known about TMS-elicited movement trajectories during the same preparatory interval. While the two types of TMS-derived measurements are likely related, e.g. movement magnitude is expected to scale with MEP amplitude, the direction of movement may be more independent of MEP amplitudes.

The ability to return to rest between task responses may hold important implications for motor system diseases that can influence the ability to modulate CSE. For example, investigations of Parkinson’s disease (Valls-Solé et al. 1994) and dystonia (Mavroudakis et al. 1995; Ikoma et al. 1996) have shown abnormal patterns of CSE at rest. Whether these out-of-task abnormalities impact patients’ abilities to return back to a resting state within a task context may help to explain specific behavioral deficits and potentially track disease progression or point toward candidate biomarkers. Examining recruitment curves in these populations during resting epochs between trials of a task may also reveal properties of the motor system responsible for the resumption of rest.

A limitation of the current study was the lack of precise feedback to participants in the process of returning to rest between task trials. EMG activity was monitored throughout the task, and the experimenter instructed participants to relax their muscles if the EMG showed signs of muscle contraction and, thus, participants could have used a variety of strategies to relax their muscle activity (e.g. ruminating on past errors or making predictions about future responses). Nevertheless, despite the unconstrained nature of this task epoch, the results suggest this inter-trial period yields a consistent pattern of CSE. Fatigue may have also limited CSE modulation due to the long duration of the experiment (Kotan et al. 2015; Morris & Christie 2020). However, the order of the hand and finger choice tasks was counterbalanced to partially control for fatigue effects. Further research under a wider variety of task conditions would be useful for determining whether the current findings generalize beyond the hand and finger choice tasks used here.

In conclusion, our current data indicate healthy young adults can promptly return to a resting CSE state within seconds following a motor response, indicating rest within the motor system is relatively stable outside and inside motor tasks. Moreover, our findings support the use of inter-trial epochs as a reference for active behavioral states, which is a common practice in physiological investigations of motor system dynamics. Our approach may have specific utility in the context of motor disorders for determining whether deficits in motor function arise from difficulty maintaining and returning to a stable resting state.

## Author Contributions

All authors contributed to the study conception and design. Material preparation, data collection and analysis were performed by Kate Bakken, Chris Horton, Mitchell Fisher, Ian Greenhouse, Charlie Lewkowitz, Hayami Nishio, and Tania Sarabia. The first draft of the manuscript was written by Kate Bakken and all authors commented on previous versions of the manuscript. All authors read and approved the final manuscript.

